# A Quantitative Design Guideline for Biomolecular Positive Feedback Systems

**DOI:** 10.1101/2025.11.09.687517

**Authors:** Vinod Kumar, Shaunak Sen

## Abstract

Feedback is at the core of biological systems found in medicine and in biotechnology. While design metaphors from control engineering are widely used to understand negative feedback in such systems, they are relatively uncommon for positive feedback, especially for biomolecular circuits. Here, we extended a block diagram modelling framework for the design of positive feedback. We found a quantitative design guideline for the strength of the positive feedback, which when wrapped around a saturation function can give a threshold and a hysteretic response. The critical feedback strength was inversely proportional to the saturation value and directly proportional to the input scale where saturation starts. We found that this saturation-threshold-hysteresis hierarchy persisted in a realistic model of a positively autoregulated gene. We showed how this classical model fitted well in a block diagram framework with multiplicative feedback and derived expressions for the critical feedback strength in terms of the saturation parameters. The dependence of the critical feedback on the parameters matched with the obtained design guideline. To complete a rigorous workflow, we discussed how Groebner Bases computations and an Interval Newton algorithm can be used to provide validated numerical solutions in biological positive feedback systems. These results should be helpful in the analysis and design of biomolecular systems with applications to the control of biomedical systems and in biotechnology.

## I. Introduction

Feedback control, both negative and positive, is extremely common in biology ([1], [2], [3]). The use of negative feedback in biology largely parallels its role in control engineering. For example, negative feedback underlies robust adaptation and homeostasis in biology, which is analogous to the integral control property of reducing the steady-state error to zero ([4]). A similar perspective has been noted for the diagnosis and treatment of diseases ([5]). The use of positive feedback, however, can be quite different. Positive feedback is typically avoided in most control engineering contexts because of possible instability. It is this same instability, however, that is harnessed in biological contexts to create memory and make decisions. A block diagram model contrasting the opposite functional roles of negative feedback and positive feedback is shown in Figure 1a ([6]): Unity negative feedback converts a large gain to one. Unity positive feedback converts a unit gain to a large gain. While block diagram models are well-developed for negative feedback design, they are relatively uncommon for positive feedback design, especially in biomolecular contexts.

**Fig. 1:**
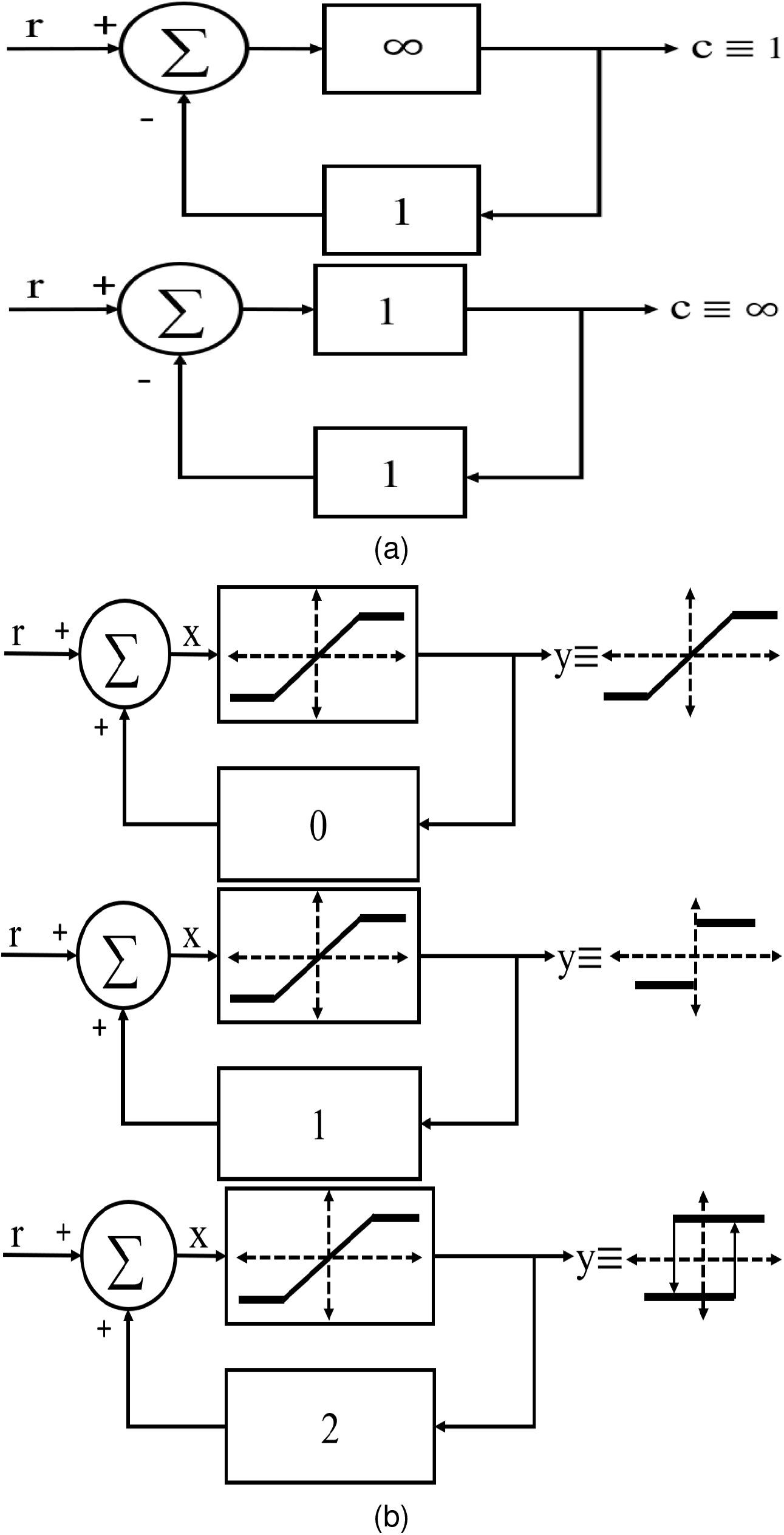
Block diagram models (a) opposite functional roles of negative feedback and positive feedback, (b) effect of the positive feedback strength (0 or 1 or 2) on the overall close-loop system response when the open-loop response is a saturation function.

Positive feedback plays an important role in the analysis and the design of biological processes at a variety of scales, ranging from viruses to cells and from cells to organisms. One of the most well-characterized systems with positive feedback in biology is the maturation of *Xenopus* oocytes ([7]). The system converts an input stimulus into an all-or-none decision representing maturation of the oocyte. A combination of mathematical modeling and experimental measurements has led to a picture where, depending on the strength of the positive feedback, the response can be switch-like or hysteretic. Other important examples of a cell-fate decision are in the choice between lysis and lysogeny by the bacteriophage *λ* ([8], [9]) and in the choice between viral replication and latency in HIV infected cells (see [10] and references therein). Further, synthetic positive autoregulatory circuits have been designed to demonstrate multiple steady-states in *E. coli* ([11]) and in mammalian cells ([12]). Finally, bistability is believed to underlie important biomedical processes such as tissue repair ([13]) and Major depressive disorder ([14]).

Sophisticated mathematical models based on nonlinear dynamical systems and differential equations are an important component of investigations into the role of positive feedback in biology. However, detailed mathematical analysis of such systems is often computationally intensive and involves numerical approximations. Strikingly, a block diagram model of positive feedback wrapping a saturation function succinctly captures the saturation-threshold-hysteresis hierarchy as the feedback gain is increased (Figure 1b, [6]). Given the successful use of block diagram models for negative feedback design, it is desirable to develop similar models for positive feedback design. In particular, a quantitative version of this hierarchy, clearly delineating the relationship between the feedback gain and the parameters of the saturation function, may organize the understanding of both analysis and design goals in a systems-level manner.

We asked how the block diagram model of positive feedback can be quantitatively extended to provide design guidelines in biomolecular contexts. We found that the critical positive feedback gain for threshold and hysteresis is related to the parameters of the saturation function: gain is inversely proportional to the saturation value and directly proportional to the input level where the saturation starts to occur. We used the classical model of a positively autoregulated gene to show that this saturation-threshold-hysteresis hierarchy persists in realistic biomolecular contexts as the effective positive feedback is increased. We showed how the critical positive feedback gain depends on the parameters of the saturation function in a manner exactly similar to that obtained from the block diagram model. We discussed how Groebner Basis computations and an Interval Analysis algorithm can help generate rigorous solutions for this model as well as for the benchmark *Xenopus* model. These results provide a quantitative systems-level design guideline for the saturation-threshold-hysteresis hierarchy in biomolecular positive feedback systems, as well as a discussion of methods that can contribute to their rigorous analysis and design.

## II. Methods

This section briefly describes the methods and the benchmark model used in this study. The methods include the computation of Groebner Bases and the basics of Interval Analysis. These methods can facilitate rigorous solutions in biomolecular contexts. Additionally, a simplified mathematical model for the effect of positive feedback in the *Xenopus* oocytes is described.

### A. Groebner Bases Computations

A Groebner Basis is an algorithmically useful generating set for an ideal of multivariate polynomials ([15]). Using Groebner Basis computations ([16]), a system of multivariate polynomial equations, which might represent a system in biology, physics, or engineering, is transformed into an equivalent form. The structure of this special equivalent form (basis) is simpler, which makes it easier to analyze the system’s properties, such as solving for roots, eliminating variables, computing dimensions/degrees, and finding multiplicities.

The main advantage of the Groebner Bases computation is that it transforms a “hard” problem into an “easy” one that is algorithmically solvable. Unlike numerical methods like the Newton’s method, which approximates solutions, Groebner bases can provide exact algebraic solutions. This method offers a well-defined procedure, via the Buchberger’s algorithm, to determine desired properties of the system ([17]). For example, the method can be used to determine whether a system has any solutions, a finite number of solutions, or infinitely many solutions. The key characteristic is the ability to eliminate variables systematically, which is extremely useful for simplifying a complex model ([18], [19]) and for understanding the relationship between key variables.

The ability of Groebner’s analysis has been leveraged in recent studies to generate insights into multi-stationarity ([20], [21]), bistability ([22]), and switchability phenomena, which are crucial for understanding and engineering biomolecular circuits. Thus, as the increasing precision of algebraic and symbolic computation widens the path from approximate numerical exploration to exact analytical characterization, Groebner Bases computations have the potential to emerge as a powerful method for analysis and design in biomolecular circuits. The starting point for the computations is the ideal ‘I’ that contains the set of polynomials for which the steady-state solutions have to be obtained. In SageMath ([23]), a standard set of statements is used to define a polynomial ring of involved variables, then an ideal set ‘I’ is defined over that ring. The built-in Groebner Basis function applies Buchberger’s algorithm to generate the Groebner Bases. The computational codes are in the GitHub repository ([24]).

### B. Interval Newton Algorithm

Interval analysis (for example, see [25], [26]) offers rigorous and practical methods that are a natural fit to the above mathematical models (see [27] for an example). These are naturally suited to the above problems because most parameters and outputs are rarely known exactly. Indeed, the fundamental nature of measurement is to provide an interval rather than a fixed value. The methods are rigorous in the sense that computations with intervals, while more complex than those with points, are rigorous numerical constructions that do not suffer from errors due to roundoff and finite computing precision. Finally, methods of interval analysis are practically oriented to provide a solution to steady-state values in a systematic and algorithmic way. These features are indispensable in the analysis and design of multiple steady-state systems, such as those in biology with positive feedback.

The Newton method is a classical method to numerically solve nonlinear systems of equations *f* (*x*) = 0 via the iterations *x*_*k*+1_ = *x*_*k*_ − *f* (*x*_*k*_)*/f*_0_(*x*_*k*_). In its standard point-based incarnation, it needs a good initial guess for the iteration to converge to the desired solution. In case multiple solutions exist, it is possible that the iterations oscillate between the solutions. The Interval version of the Newton method has a similar form *X*_*k*+1_ = *N* (*X*_*k*_) ∩ *X*_*k*_ where *N* (*X*_*k*_) = *mid*(*X*_*k*_) − *F* (*mid*(*X*_*k*_))*/F*_0_(*X*_*k*_), except that it is guaranteed to enclose all roots in a given starting interval *X*_0_. Further, it has many useful properties, such as the possibility of providing a criterion for the non-existence of a solution.

The interval notions of arithmetic and functions are needed to implement the Interval Newton algorithm. The uppercase letter *X* denotes intervals on the real line, whereas the corresponding lowercase letter *x* denotes a real number. Similarly, the uppercase letter *F* denotes the natural interval extension of a real-valued function *f* obtained by replacing each element by its corresponding interval. Arithmetic operations on intervals are defined in a set-theoretic sense. Division is performed using an extended interval arithmetic that allows division by intervals containing the element 0. This is the crucial step in capturing all solutions in the given starting interval. The symbol *mid* refers to the mid-point of the interval.

These computations were performed in Julia using the Interval Arithmetic package ([28]). The computer code is available on a GitHub repository ([24]).

### C. Model

Positive feedback has been found to play a key role in how *Xenopus* oocytes determine their fate ([7]). The induction of cell fate in *Xenopus* oocytes necessitates a conversion of a chemical signal (progesterone) into an all-or-none decision. Essentially, the maturation process “remembers” a transient input. A combination of experimental work and mathematical models has highlighted the role of nonlinear positive feedback in implementing such a decision. A simplified model for this process is

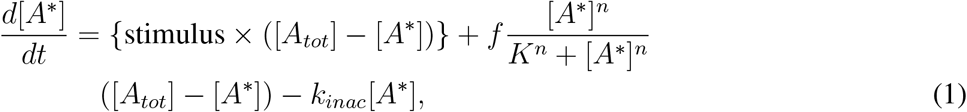

where [*A*^∗^] is the concentration of the output biomolecule, [*A*_*tot*_] is the total concentration, stimulus is the input, *f* is the feedback strength, *n* and *K* are other parameters of the feedback, and *k*_*inac*_ is a parameter for the degradation of the output. At steady-state, the right-hand side is set to zero, and the value of [*A*^∗^] is calculated for a fixed value of other parameters. A plot of this value for different stimulus values gives the input-output response. This shape has been shown to be sigmoidal for small feedback strengths and hysteretic for larger feedback strengths. The hysteresis is believed to underlie the irreversibility of the cell-fate decision.

## III. Results and Discussion

### A. Quantitative Design Guideline

A block diagram model provides a simple way to quantitatively understand the saturation-threshold-hysteresis hierarchy. A saturation function captures the essential bounded nature in most biological systems. Such bounds arise, for example, due to finite binding capacity or resource limitations in enzyme kinetics.

We considered a generic piecewise linear saturation function

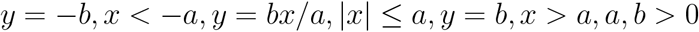

wrapped in a positive feedback loop (Figure 2a). To obtain a quantitative guideline for positive feedback, we derived a relation between the critical positive feedback gain needed for threshold and hysteresis and the parameters of the saturation function.

#### Proposition 1

For the block diagram model shown in Figure 2a, the overall closed-loop function remains a saturation function for *K* < *a/b*, is a threshold for *K* = *a/b*, and becomes hysteretic for *K* > *a/b*.

#### Proof

Because of the positive feedback, the effective input to the saturation function is *x* = *r* + *Ky*. Consider the response across each zone of the saturation function with input *x* = *r* + *Ky* and output *y*. If *r* + *Ky* < −*a*, then *y* = −*b* ⇒ *r* < −*a* + *Kb*. Similarly, if *r* + *Ky* > *a*, then *y* = +*b* ⇒ *r* > *a* − *Kb*. Finally, 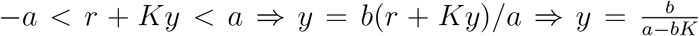 whenever min{−*a* + *Kb, a* − *Kb*} < *r* < max{−*a* + *Kb, a* − *Kb*} . Therefore, the overall closed loop function is a saturation function for 0 < *K* < *a/b*, a threshold function for *K* = *a/b*, and a hysteresis function for *K* > *a/b* (Figure 2b). □

**Fig. 2:**
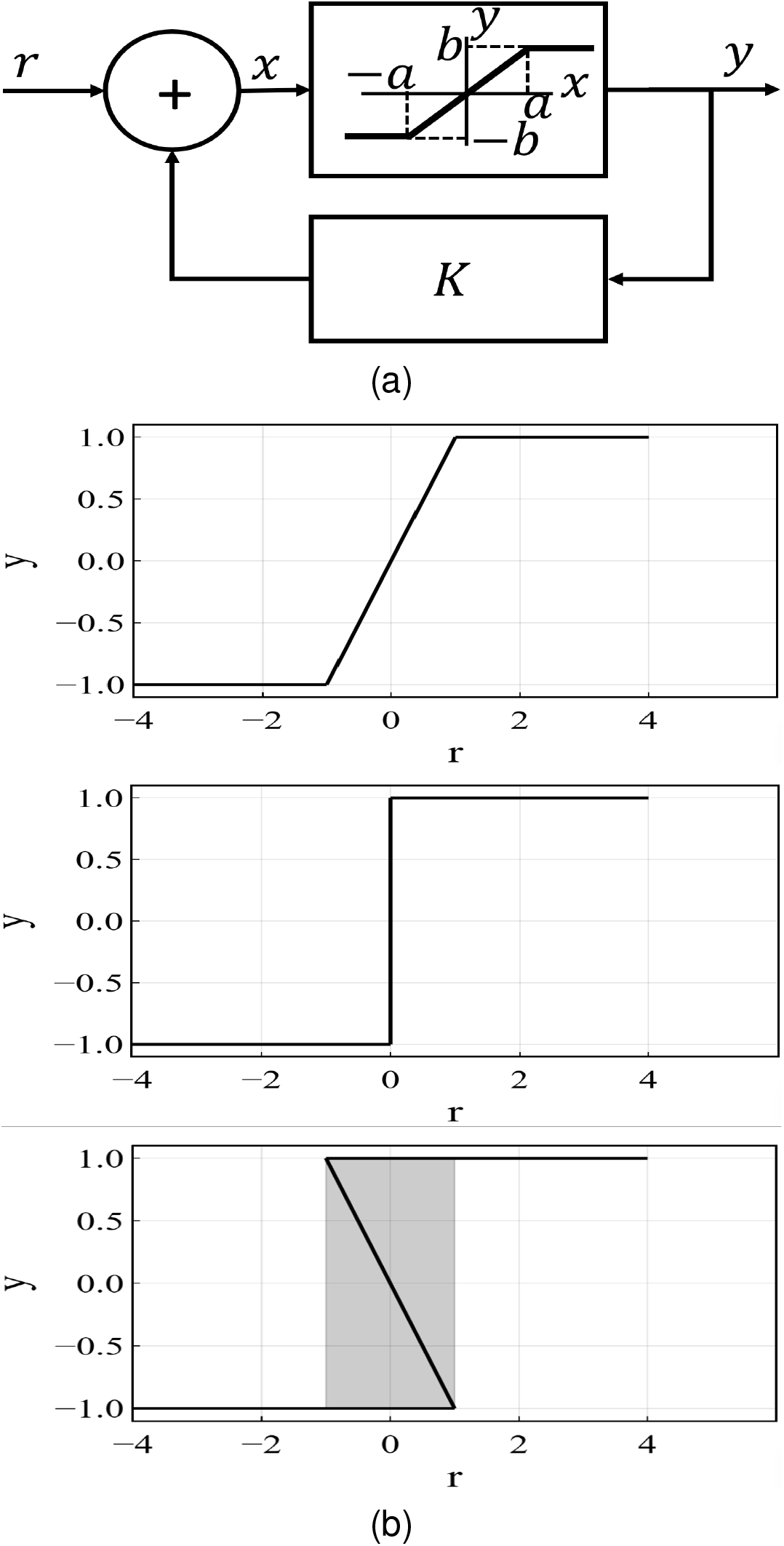
Quantitative effect of positive feedback strength. (a) A saturation function wrapped in positive feedback. (b) Closed-loop response is like a saturation function for 0 < *K* < *a/b*, a threshold function at *K* = *a/b*, and a hysteresis function for *K* > *a/b*. The shaded region highlights the hysteretic zone.

The critical positive feedback gain is *K*_*c*_ = *a/b*, where *a* represents the input value where the saturation function saturates and *b* represents the saturation output value. Therefore, depending on the strength of the positive feedback, the saturation function may remain a saturation function or transform into an overall threshold or even a hysteretic function. This block diagram model provides a simple way to quantitatively understand the transitions from a saturation function to a threshold function and from a threshold function to a hysteresis function. This is consistent with biological studies where increasing the positive feedback strength causes a transition to hysteresis. The transitions from saturation to threshold to hysteresis as feedback gain is increased were already known ([6], Figure 1a). To the best of our knowledge, however, a quantitative version had not been reported. These results add a quantitative understanding of the critical positive feedback strength in terms of the parameters of the saturation function, specifically the ratio of the input level that causes the saturation (*a*) to the overall saturation level (*b*).

### B. Positive Autoregulation Model

The block diagram model above provided key design insight into the role of positive feedback gain in determining threshold and hysteretic responses starting from a saturation function. However, realistic biomolecular circuit models based on mass action kinetics differ from the block diagram model in at least three crucial ways. First, the saturation function is usually differentiable and not piecewise linear. Second, the biomolecular concentrations are positive and not symmetric about zero. Third, the positive feedback is implemented using biomolecular interactions and not just through a gain block. To check the persistence of the design guideline *K* > *a/b* in these contexts, we analyzed a simple model of a positive autoregulatory gene.

A reduced model for such a positive autoregulatory circuit is commonly used to show how positive feedback can generate hysteresis ([3]). The core of the circuit has a gene that expresses a transcription factor. The transcription factor can dimerize and bind to the promoter of the gene. When the transcription factor is bound to the promoter, RNA polymerase binds to the promoter, and more copies of the transcription factor are generated. Therefore, the presence of a transcription factor results in more copies of the transcription factor, giving rise to positive feedback. The mathematical model of the positive autoregulation circuit that we derived emphasizes the input and the output as well as the key intermediate steps, so that the correspondence with the above block diagram framework is clearer.

First, we considered the origin of saturation in the model of the positively autoregulated gene. Consider the Model S, where the RNA Polymerase *R* is the input that activates transcription by binding to DNA *D*’s promoter region (inset Figure 3a ). This produces the protein output *P* . The map from the input (*R*) to the output (*P* ) can be obtained as follows: DNA *D* is assumed to exist in two states — an inactive state *D*_*U*_ (unbound DNA) that does not produce *P* and an active state *D*_*R*_ (RNA polymerase-bound DNA) that does produce *P* (inset Figure 3a). The conversion between these two states follows the reactions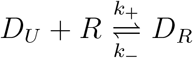, where *k*_+_ and *k* denote the first-order association and dissociation rates, respectively. Since the total DNA is conserved, the conservation law *D*_*U*_ + *D*_*R*_ = *D*, where *D* is the total DNA concentration, holds. At steady-state, the forward and reverse reaction rates are equal *k*_+_*RD*_*U*_ = *k*_−_*D*_*R*_. Using *D*_*U*_ = *D* − *D*_*R*_, we get,

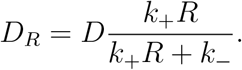

**Fig. 3:**
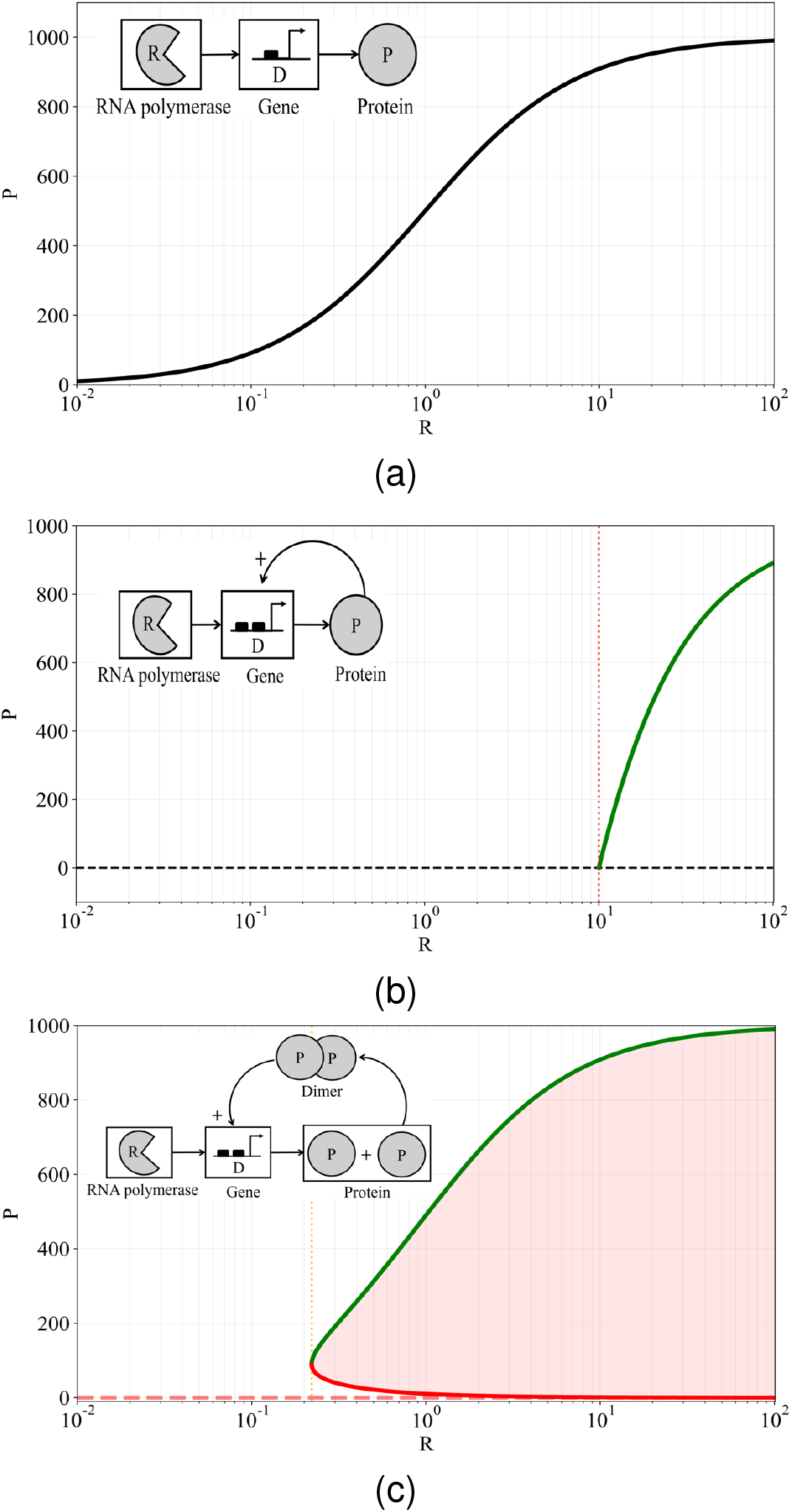
Input-output responses for (a) Model S, showing saturation, (b) Model T, showing threshold (red dashed line), and (c) Model H, showing hysteresis (red shaded region). Insets show a schematic of the corresponding biomolecular circuits. Parameters: *D* = 10, *k*_+_ = 100, *k*_−_ = 100, *k*_1+_ = 100, *k*_1−_ = 100, *α* = 100 and *γ* = 1, *k*_2+_ = 0.1, and *k*_2−_ = 1000.

The reactions for the production of the protein *P* from the active DNA state *D*_*R*_ and its degradation via a first-order reaction with rate constant *γ* are 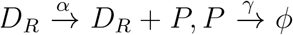 . At steady-state, *αD*_*R*_= *γP* . Therefore, the input-output response is

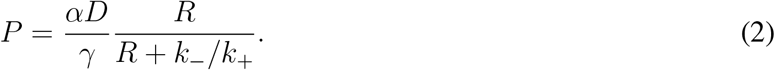

This input-output response is an instance of a saturation function with *a*∼ *k*_−_*/k*_+_ and *b* ∼ *αD/γ* (Figure 3a).

Second, we added a positive feedback through the requirement that the RNA polymerase binds *R* to the promoter only when the transcription factor protein *P* is already bound to the promoter (Model T in inset Figure 3b). DNA *D* is assumed to exist in three states — two inactive states *D*_*U*_ and *D*_*P*_ (protein-bound DNA) and an active state *D*_*PR*_ (RNA polymerase-bound DNA) that does produce *P* . The reactions for the binding-unbinding of *P* to *D*_*U*_ and *R* to *D*_*P*_ are 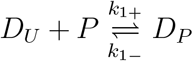 and 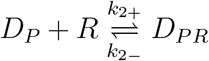, respectively. The association and dissociation rate constants are noted above and below the respective reaction directions. The conservation law gives *D*_*U*_ + *D*_*P*_ + *D*_*PR*_ = *D*. The production of the protein from the state *D*_*PR*_ and its degradation proceeds as: [ineq (inset Figure 3b). At steady-state, *k*_1+_*PD*_*U*_ = *k*_1−_*D*_*P*_, *k*_2+_*RD*_*P*_ = *k*_2−_*D*_*PR*_, and *αD*_*PR*_ = *γP* . Solving for the input-output response necessitates algebraic manipulations. From the steady-state,

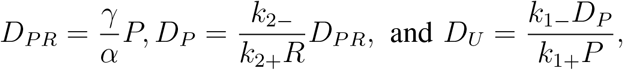

where the last equation holds when *P*≠ 0. If *P* = 0, then *D*_*U*_ = *D, D*_*P*_ = *D*_*PR*_ = 0 is a solution. If *P*≠ 0, the conservation law gives

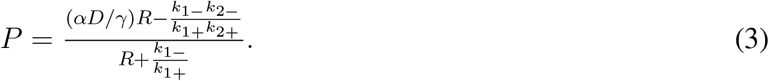

This solution exists whenever it is positive, which is when the input 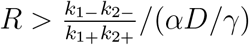. The input-output response is shown in Figure 3b.

Third, we considered a model where the dimerization of the output protein is needed for it to bind to the DNA and activate expression (Model H in inset Figure 3c). The requirement of such co-operativity adds another layer of positive feedback to the model. The input-output response is obtained using the reaction assumptions and conservation law in a manner similar to those computed in the previous cases.

The reactions are 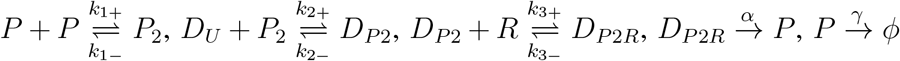 and 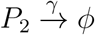 the conservation law is *D*_*U*_ + *D*_*P* 2_ + *D*_*P* 2*R*_ = *D*. At steady-state,

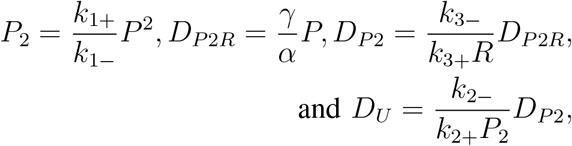

if *P*_2_ ≠ 0, or equivalently *P* ≠ 0. If *P* = 0, *D*_*U*_ = *D* and *D*_*P* 2*R*_ = *D*_*P* 2_ = 0. If *P* ≠ 0, the conservation law can be used to get the quadratic equation.

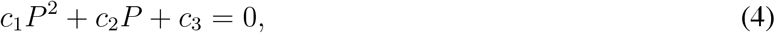

where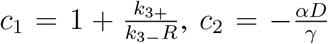, and 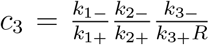. When the discriminant 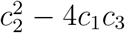 is positive, there is a possibility of hysteresis in this system. Therefore, this additional positive feedback due to dimerization can give a hysteretic response (Figure 3c). We have shown how the positive feedback in a positively autoregulatory model can also show the transition from saturation to threshold and from threshold to hysteresis. We presented a detailed derivation of a relatively standard model to emphasize the connections to the block diagram model. The simplest choice of reaction assumptions that demonstrate the connections was used. We concluded that the results seen in the block diagram model have analogs in realistic contexts as well.

### C. Multiplicative Feedback Block Diagram Model

Typically, analysis of Model H and similar models relies on a bifurcation theory framework and associated numerical simulations to chart out the region in parameter space where bistability can occur. This does not, by itself, decouple into a form where the critical feedback strength is expressed in terms of other parameters. Given the *K* > *a/b* relation derived above, which explicitly links the critical feedback strength to the parameters of the saturation function, we investigated whether a similar block diagram representation and interpretation for critical feedback strength are possible.

We found that the input-output responses for Models S, T, and H could be represented in a block diagram model with multiplicative feedback, and where the blocks represented nonlinear blocks (Figure 4). The forward path is the saturation function *P* = *s*(*R*), where 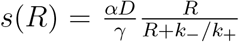 . This corresponds to *a*∈ *k*_−_*/k*_+_ and *b* ∈ *αD/γ*. For Model T and Model H, there is also a feedback path 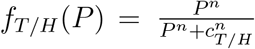, with *n* = 1 and *n* = 2, respectively. For Model T, *c*_*T*_ = *k*_1−_*/k*_1+_ and 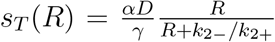, *a*_*T*_ = *k*_2−_*/k*_2+_, *b*_*T*_ = *αD/γ*. For Model H, *c*_*H*_ = (*k*_2−_*/k*_2+_)((*k*_1−_ + *γ*)*/k*_1+_) and 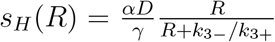, *a*_*H*_ = *k*_3−_*/k*_3+_, *b*_*H*_ = *αD/γ*. The overall closed-loop feedback relation is *P* = *s*(*Rf* (*P* )). For fixed *n*, the feedback strength is proportional to 1*/c* as a small value of *c* ensures the feedback is activated for small values of *P* and vice versa. Therefore, the feedback strengths for Model T and Model H are taken as *K*_*T*_ = 1*/c*_*T*_ and *K*_*H*_ = 1*/c*_*H*_, respectively.

**Fig. 4:**
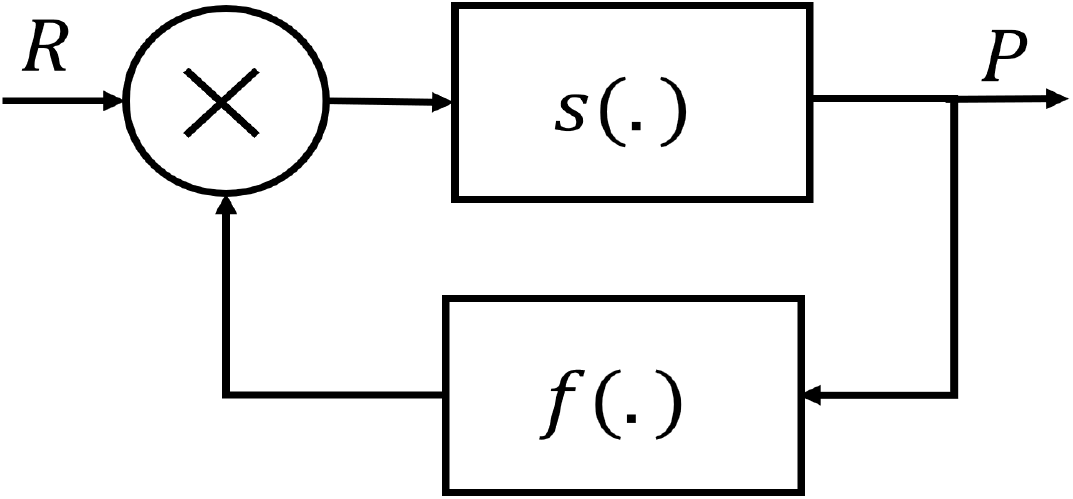
Block diagram model with multiplicative feedback.

We summarized the analogous quantitative guidelines for Model T and Model H in Proposition 2 and Proposition 3, respectively.

#### Proposition 2

For the block diagram model shown in Figure 4 with *n* = 1 (Model T), the overall closed-loop function is a threshold for *K*_*T*_ > *a*_*T*_ */b*_*T*_ *R*.

#### Proof

The proof follows from *P* = *s*_*T*_ (*Rf*_*T*_ (*P* )). *P* = 0 is clearly a solution. If *P* ≠ 0, *P* = (*b*_*T*_ *R − a*_*T*_ *c*_*T*_ )*/*(*R* + *a*_*T*_ ), which exists when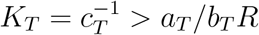, as desired. □

#### Proposition 3

For the block diagram model shown in Figure 4 with *n* = 2 (Model H), the overall closed-loop function is a hysteresis for 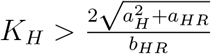

#### Proof

The proof follows from *P* = *s*_*H*_ (*Rf*_*H*_ (*P* )). *P* = 0 is clearly a solution. If 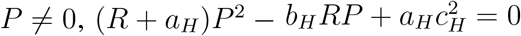 . Two solutions exist if the discriminant 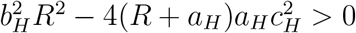. This gives the desired condition 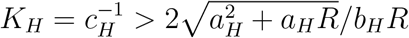, as desired. □

#### Remark 1

The condition obtained in Proposition 3 is not of the form *K* > *a/b*. However, it preserves the proportionality relation that the critical feedback strength is a ratio in which the numerator and denominator are increasing functions of *a* and *b*, respectively.

#### Remark 2

The critical feedback strength in Proposition 2 and Proposition 3 depends on the input strength *R*, unlike the case with Proposition 1. This is possibly because of the multiplicative nature of the feedback. We noted that the quantitative guidelines derived in Proposition 2 and Proposition 3 are of great value. While these are mere rearrangements of the steady state equations derived in the earlier subsection, their form clearly delineates the feedback strength from the parameters of the saturation function. That the structure is of the same form as obtained in the subsection on the block diagram model, lends credence to the persistence of the core message *K* > *a/b*, for at least a class of systems.

## IV. Discussion

The saturation-threshold-hysteresis hierarchy is present in diverse engineering and biological contexts with positive feedback. Here, we developed a block diagram model to quantitatively map how this hierarchy depends on the positive feedback gain and the parameters of the saturation function that is enveloped by the feedback. We used a concrete example of a gene that activates itself, showing how successive enhancement of the positive feedback follows this hierarchy. Further, we showed how this realistic model generates a quantitative design guideline for threshold and hysteresis that is very similar to the one obtained from the block diagram model. Finally, we discussed below the methods that can be used for a rigorous analysis of such systems, illustrating them with the example of the positively autoregulated gene model, as well as a classical Xenopus model. These results should contribute to the analysis and design of biomolecular positive feedback circuits, such as for memory and decision-making.

The simple model derived above was solved to obtain analytic closed-form solutions, and numerical simulations were used to plot these solutions. Typically, such closed-form analytical solutions cannot be obtained, even for simple variants of the above model. However, there are methods that can be used to obtain rigorous solutions. These augment typically used geometric approaches that rely on the existence of one or more points of intersection between two curves with concrete solution values. We discussed two such methods based on the computation of a Groebner Basis ([29]) and Interval Analysis ([25], [26]).

### A. Groebner Basis Computations

The computation of a Groebner Basis provides a strategy to automate the algebraic manipulation used to obtain solutions in the above model. The steady-state equations obtained in the above models constitute a set of polynomial, not linear, equations. Just like Gaussian Elimination for linear equations, Groebner Basis methods provide a way to simplify these polynomial equations.

We used SageMath version 9.3 for doing the Groebner Basis computations and the other analytical simplifications. For computing the steady-states of the models, the equations were converted to systems of polynomial equations. The complete set of equations consists of binding, conservation, and protein input-output relations. The lexicographic ordering used for computing the Groebner Basis was *D*_*U*_ > *D*_*R*_ > *P* > *R, D* > *D*_*U*_ > *D*_*P*_ > *D*_*PR*_ > *P* > *R*, and *D*_*U*_ > *D*_*P* 2_ > *D*_*P* 2*R*_ > *P*, respectively.Table I displays the Groebner Bases for generic parameter values to illustrate the general dependence on parameters. The equations in the Groebner Basis lend themselves to further exact or numerical solutions. We considered the following parameters for the numerical computations, *D* = 10, *k*_+_ = 100, *k*_−_ = 100, *k*_1+_ = 100, *k*_1−_ = 100, *α* = 100 and *γ* = 1, *k*_2+_ = 0.1, and *k*_2−_ = 1000. *D* has a unit of concentration (*nM* ), and the rate constants are in *M* ^1−*n*^*S*^−1^, where *n* is the overall reaction order with respect to the rate constant in question.

**TABLE 1:**
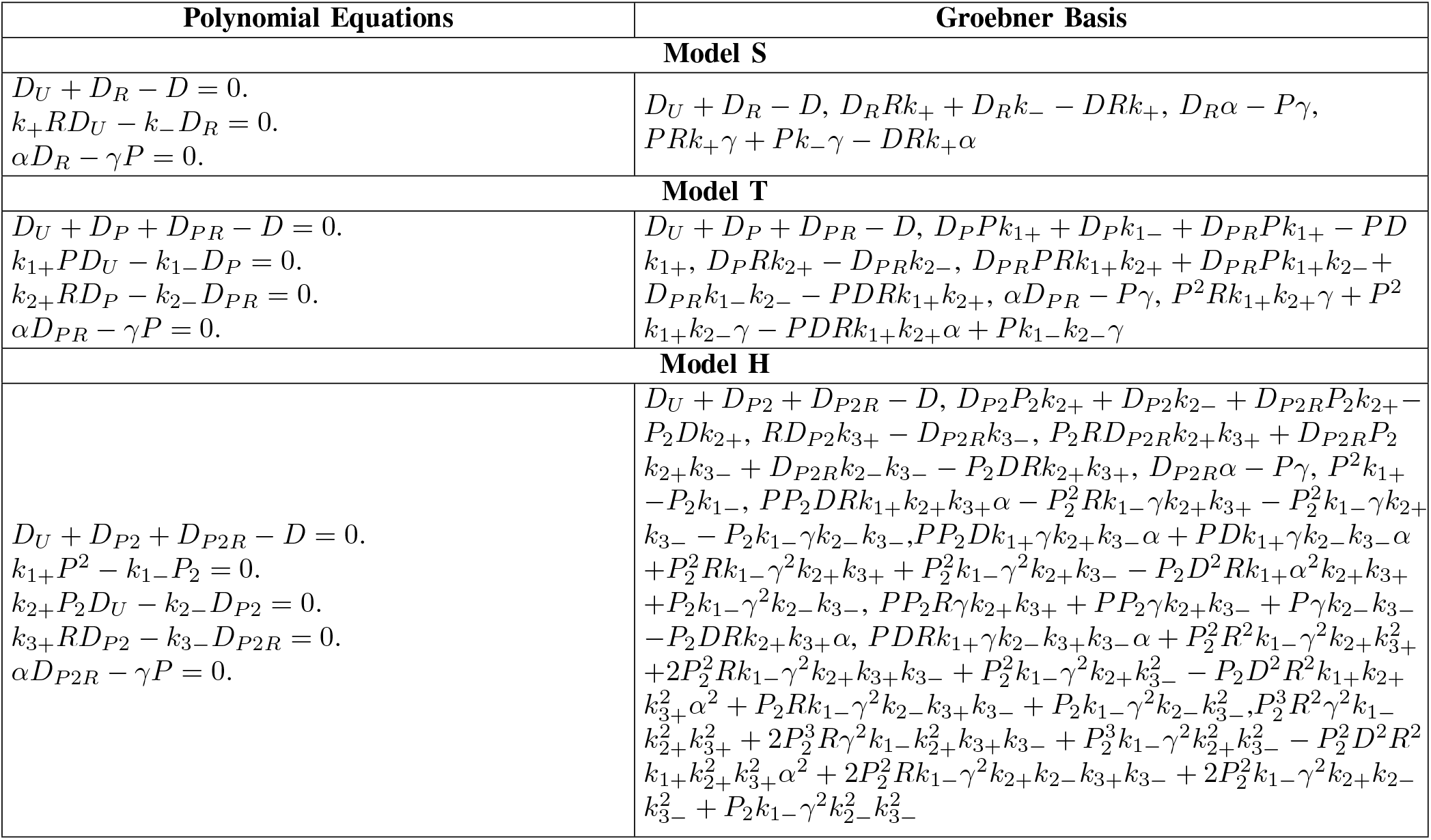
Polynomial equations and Groebner Basis for different cases.

### B. Interval Newton Algorithm

We showed how the benchmark *Xenopus* model can be rigorously solved using the Interval Newton algorithm. The steady-state equation is given in Equation (1). Obtaining analytical closed-form solutions for the steady-states are challenging. Therefore, numerical or geometrical methods are typically used. These have shown how the response as *f* changes moves from sigmoidal to hysteretic ([7]). The parameter used were *f* = (0 to 1.4)*/*10, *k*_*inac*_ = 1, *n* = 5, *A*_*tot*_ = 1, *K* = 1, stimulus = −2.5 to −1.5. For a specific value of the parameter set (*k*_*inac*_ = 0.01, *n* = 5, *A*_*tot*_ = 1, *K* = 1), geometric methods were used to find the intersection between two curves whose solutions give the steady-state for each value of the stimulus. The red line is the stimulus. Blue line is the plot between *A*^∗^*/A*_*tot*_ and 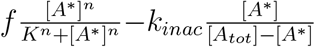 (Figure 5). The computations, however, are approximate because of approximations in the floating-point representation. The Interval Newton algorithm can be used with an initial guess of [0, 1] (as the steady-state is guaranteed to lie between these two limits from biological constraints). The algorithm guarantees a rigorous bound on all solutions in this interval. For *f* = 0.14, the steady-states are guaranteed to lie in the intervals

**Fig. 5:**
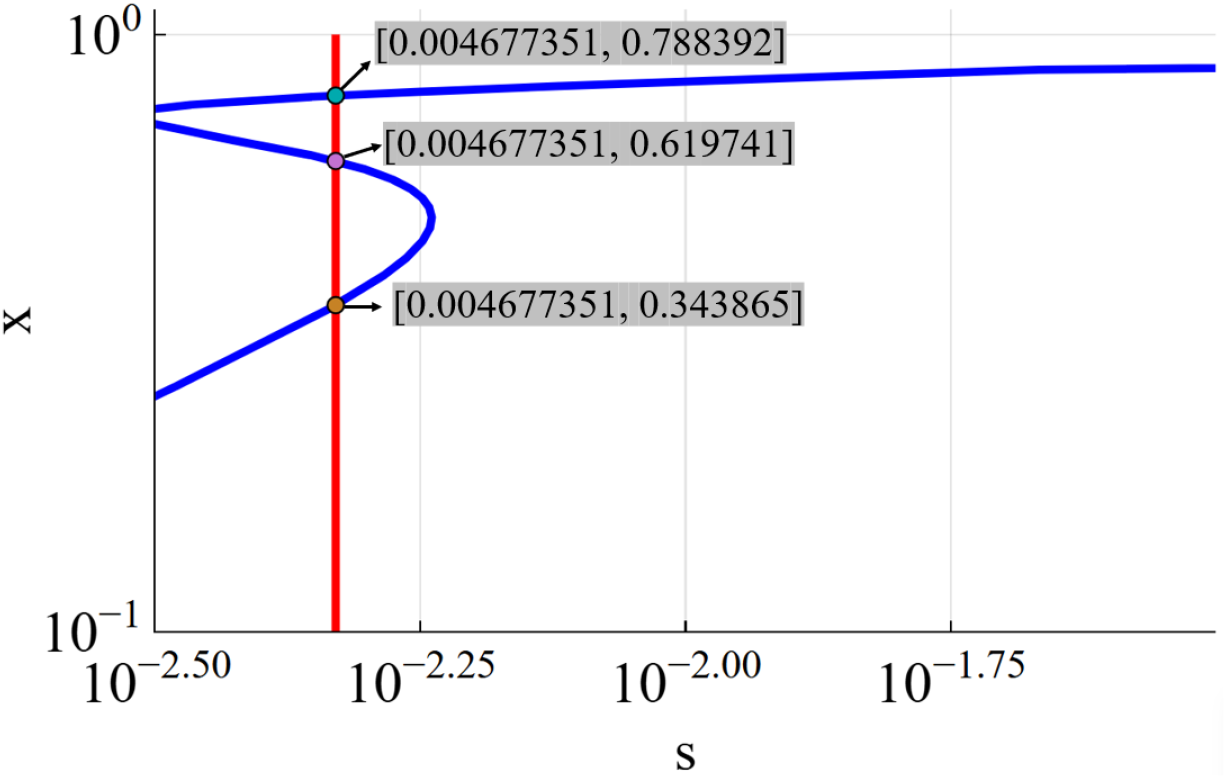
Steady-state solution in the *Xenopus* model. For the blue line, the Y-axis represented by X is *A*^∗^*/A*_*tot*_, and the x-axis is 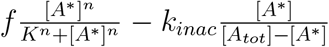 . The red line is just the value of the stimulus on the X-axis. Indicated points are the solutions obtained by the Interval Newton algorithm.

[0.61974, 0.619741], [0.788391, 0.788394], and [0.343855, 0.343872] (dots in Figure 5). These compare very well with the numerically obtained approximate solution. The advantages of this approach are that the computation represents a rigorous numerical construction and all solutions are found directly.

## V. Conclusion

Decision-making and memory are important systems-level phenomena prevalent in diverse biological contexts. The underlying principle has been understood to be positive feedback. However, unlike the situation for negative feedback, quantitative systems-level design guidelines for positive feedback do not exist. We showed how simple block diagram models can be used to generate these design guidelines and relate the critical feedback strength for threshold and hysteresis to the key parameters of the saturation function that is inside the feedback loop. These guidelines seamlessly transition to a realistic model of a positively autoregulated gene. Additionally, we discussed how methods from algebraic geometry and Interval analysis can help in rigorously predicting multi-stability in a wide range of biomolecular circuits, complementing the exhaustive, but approximate numerical simulations. Overall, this work should facilitate efforts for the analysis and design of positive feedback in biological contexts with potential for applications in biomedical and biotechnological control.

